# Deciphering the combinatorial influence of diet and the microbiota on experimental colitis

**DOI:** 10.1101/117929

**Authors:** Sean R. Llewellyn, Graham J. Britton, Eduardo J. Contijoch, Arthur Mortha, Jean-Frederic Colombel, Ari Grinspan, Jose C. Clemente, Miriam Merad, Jeremiah J. Faith

## Abstract

**Background & Aims:** The complex interactions between diet and the microbiota that influence mucosal inflammation and inflammatory bowel disease are poorly understood. Experimental colitis models provide the opportunity to control and systematically perturb diet and the microbiota in parallel to quantify the contributions between multiple dietary ingredients and the microbiota on host physiology and colitis.

**Methods:** To examine the interplay of diet and the gut microbiota on host health and colitis, we fed over 40 different diets with varied macronutrient sources and concentrations to specific pathogen free or germ free mice either in the context of healthy, unchallenged animals or colitis models (dextran sodium sulfate (DSS) and T cell transfer).

**Results:** Diet influenced physiology in both health and colitis across all models, with the concentration of protein and psyllium fiber having the most profound effects. Increasing dietary protein elevated gut microbial density and worsened DSS colitis severity. Depleting gut microbial density by using germ-free animals or antibiotics negated the effect of a high protein diet. Psyllium fiber influenced host physiology and attenuated colitis severity through microbiota-dependent and microbiota-independent mechanisms. Combinatorial perturbations to dietary protein and psyllium fiber in parallel explain most variation in gut microbial density, intestinal permeability, and DSS colitis severity, and changes in one ingredient can be offset by changes in the other.

**Conclusions:** Our results demonstrate the importance of examining complex mixtures of nutrients to understand the role of diet in intestinal inflammation.

---

Despite strong patient interest and numerous studies implicating diet in the exacerbation or prevention of inflammatory bowel disease (IBD), exclusive enteral nutrition (EEN), whereby patients consume a nutritionally complete simple diet, provides the only clinically validated diet that can treat IBD, specifically pediatric Crohn’s disease (CD).^1-3^ However, the specific mechanisms driving therapeutic response to EEN are poorly understood, and the return to a normal diet is typically accompanied by the return of the colitis. Furthermore, consumption of EEN is too divergent from normal dietary customs to represent a practical maintenance therapy. The ideal IBD diet would have higher efficacy than EEN and allow the consumption of a diet with few restrictions to maximize patient compliance. Developing such a diet for IBD is a major challenge, as diet’s influence on mucosal inflammation likely involves complex interactions between multiple dietary ingredients, nutrients, host physiology, disease heterogeneity, and the gut microbiota.

Many of the microbes that colonize our gut microbiota are stably colonized for decades,^4^ but their relative abundances are in constant in flux as our gut microbial ecosystem responds to the nutrients within our diet.^5–10^ In both mice and humans, the abundances of bacterial community members can change several orders of magnitude within 24 hours of changing diet, remain stable while diet is held constant, and reverse when diet is reverted.^5–8, 11^ There are numerous lines of evidence suggesting the gut microbiota plays a role in IBD, including differences in the microbiota composition of individuals with IBD compared to healthy controls,^12,13^ the requirement of the gut microbiota for disease initiation in multiple animal colitis models,^14^ and the impact of intensive fecal microbiota transplantation in treating ulcerative colitis (UC).^15,16^ Furthermore, diet and the microbiota can work in concert to influence host physiology through both metabolites and immune interactions.^17–21^

To understand the complex interactions between multiple dietary ingredients, nutrients, and the microbiota in a controlled manner, we used specific pathogen free (SPF) and germ free (GF) animals combined with over 40 unique diets to quantify the individual and synergistic influence of dietary macronutrients and the microbiota on host physiology and experimental colitis.

## Materials and Methods

*Mice* — SPF mice were purchased from Jackson Labs (C57BL/6J mice) or bred in-house (Rag1^-/-^ C57BL/6J mice). GF mice were housed in standard flexible film isolators. Male mice were used for dextran sulfate sodium (DSS) experiments in SPF mice. All other experiments used both male and female mice.

*Diets* — Custom refined mouse chows (**Table S1**) were designed in collaboration with a nutritionist at Envigo. Carbohydrates were subdivided into host digestible carbohydates (which we will simply refer to as carbohydrates) and host indigestible carbohydrates (fiber). For each macronutrient source we preferentially choose refined sources that contain only a single macronutrient although some tested ingredients (e.g., fiber sources), contain a mixture of macronutrients. The macronutrient sources included animal and plant proteins, animal and plant fats (with diverse fatty acid profiles), simple and complex sugars, and dietary fibers. All custom diets were irradiated and vacuum-sealed.

*Dextran Sulfate Sodium Colitis Model* — One week prior to the administration of DSS, mice were provided a custom diet (**Table S1**) *ad libitum* and were weighed daily for the duration of the experiment. After a week of diet *ad libitum,* the mice were continued on the same diet and given 3% DSS in drinking water to induce mucosal injury. Mice were sacrificed at day 7 or upon reaching a humane endpoint (i.e., >18% weight loss, a body condition score of 2, or based on humane indicators). GF mice were given 2% DSS instead of 3% DSS to induce roughly equivalent survival times in SPF and GF mice (**Figure S3)**. For animals sacrificed for ethical reasons prior to day 7, we extrapolated the weight loss curve of each animal by linear regression using their last three measured time points. These extrapolated values are highlighted in yellow in **Table S2A** and as a dotted line in **Figure 2A**. In addition, for analysis of survival time the date of death/sacrifice was the time at which animals reach 18% weight loss or other humane endpoint.

*Adoptive T Cell Transfer Colitis Model* — One week prior to T cell transfer, 8-12 week-old GF or SPF Rag1^-/-^ C57BL/6J mice were given a custom diet *ad libitum* for the duration of the experiment. Each mouse received an intraperitoneal injection of 10^6^ naive T cells (CD4+CD25^-^ CD45RB^Hi^) isolated with MACS and FACS at 98% purity from age and sex matched WT C57BL/6J mice.^22^ Ten-percent of bedding was retained at each cage change to prevent fluctuations in animal environment.

*Colonic Cytokine Quantification* — Each colon had its fecal contents removed and was sliced along the mesentery. Colons were washed twice in PBS, and incubated in tissue culture media for 24 hours at 37°C. Cytokine concentrations were assayed in tissue culture supernatant with either a sandwich ELISA for IL-6 and TNF-α (Biolegend; **Fig. 2 and S1**) or a custom Milliplex Map Kit for IL-10 and TNF-α (Millipore; **Fig. 3**)

*Lipocalin-2* — To measure intestinal inflammation longitudinally,^23^ fecal pellets were weighed, homogenized in PBS, and centrifuged to remove large debris. Lipocalin-2 (R&D Systems) was measured in the supernatant by sandwich ELISA.

*Microbiota processing and analysis* — Fecal microbiota samples were processed as previously described to assay community composition and estimate microbial density.^4,24^ Each mouse fecal pellet was weighed. Fecal DNA was extracted by bead-beating and quantified with Qubit assays (Life Technologies). The 16S rDNA amplicon sequencing of the V4 hypervariable was performed with an Illumina MiSeq (paired-end 250bp). The 16S rDNA data were analyzed with MacQIIME 1.9.1.^25^ OTUs were picked with 97% sequence similarity and the sequences were aligned to the Greengenes closed reference set with a minimum sequence length of 150bp and 75% ID.^26^ LEfSe (linear discriminant analysis with effect size) was used to identify microbial clades that were significantly associated with changes in diet.^27^

*Short Chain Fatty Acid Quantification* — Short chain fatty acids were measured by gas chromatography mass spectrometry at The Stable Isotope and Metabolomics Core Facility of the Albert Einstein College of Medicine. Metabolites from cecal content samples were extracted with 0.7 ml of water containing 10 μg/ml of propanoic acid_D5, and 5 μg/ml butyric acid_D7 as internal standards. Helium was used as carrier gas at a consistent flow of 1ml/min. Chemstation was used for data analysis.

*Oral administration of Short Chain Fatty Acids in germ-free animals* — GF C57BL/6J mice were given either normal or SCFA supplemented water (67.5 mM Sodium Acetate [Sigma Aldrich], 40 mM Sodium Butyrate [Sigma Aldrich], and 25.9 mM Sodium Propionate [Sigma Aldrich]).^28^ The SCFA supplemented water was made fresh and changed weekly. After 4 weeks of diet ± SCFA water, the mice were continued on the diet *ad libitum* and given 2% DSS ± SCFA to induce a mucosal injury and test the effect of SCFA on the DSS model.

*Colonic Tissue Processing and FACS Analysis* — From each colon we removed fecal contents and mesenteric fat, sliced them along the mesentery, and washed once in PBS and twice in HBSS. Each colon was then placed into dissociation buffer (HBSS without Ca^2+^ and Mg^2+^, 10% FBS, 5mM EDTA, 15mM HEPES) on a shaker to remove the epithelium. Each colon was washed twice in HBSS and transferred into fresh pre-warmed digestion buffer (HBSS without Ca^2+^ and Mg^2+^, 2% FBS, 0.5mg/mL collagenase VIII [Sigma Aldrich], 0.5mg/mL DNase 1 [Sigma Aldrich]) and incubated on a shaker. Digested colons were vortexed briefly and ran through a 100μm strainer. The digestion and straining were repeated. HBSS was added to the suspension and centrifuged. Cells were washed once with HBSS and resuspended in PBS for staining. Cells were blocked with Fc Block (CD16/32 [Biolegend]) and stained for viability (Blue [Life Technologies]), CD4 (AF700 [Biolegend]), CD45 (APC-Cy7 [Biolegend]), and FoxP3 (PE [Biolegend]). Surface markers were stained before fixation and intracellular markers were stained after fixation with the FoxP3 Fixation/Permeabilization Kit (eBioscience). Samples were run on a BD LSRII and analyzed with FlowJo (see **Fig. S12** for gating schema).

*Intestinal Permeability* — Intestinal permeability was assessed by luminal gastrointestinal gavage of FITC-dextran (3-5kDA) [Sigma-Aldrich], a non-metabolizable macromolecule that was used as a permeability probe. Mice were fasted overnight and for the remainder of the experiment. Mice were gavaged with FITC-dextran (40mg/100g body weight) 4 hours before bleeding. The concentration of FITC-dextran in serum was quantified by spectrophotofluorometry (485/528nm).

*RNA extraction from colonic tissue and RNA-Seq* — 10-30mg of tissue from the proximal and distal colon was placed into an RNAlater (Qiagen). RNA was extracted with an RNeasy Kit (Qiagen) using 1mm zirconia/silica beads (BioSpec). mRNA Illumina libraries were generated by the NYU Genome Technology Center and sequenced to a depth of 43 ± 3 million (mean ± std) 50nt paired-end reads/sample (Illumina HiSeq). Transcript expression values were obtained by mapping to the *Mus musculus* mm10 genome (http://genome.ucsc.edu/) using HISAT^29,30^ and StringTie.^31^ Differential expression was determined with the ballgown R package.^32^

*Histology* — Samples were sectioned, stained for H&E and imaged by HistoWiz Inc. (histowiz.com). Whole slide scanning (40x) was performed on an Aperio AT2 (Leica Biosystems) and a pathologist scored the resulting images.

*Computational modeling and feature selection* — Intestinal permeability, weight change, survival, and gut microbial density were modeled as a function of dietary composition using linear regression. Lasso and stepwise-regression were used to identify the dietary factors that best explained the observed variation in host physiology or microbial density using R and JMP. Best models by stepwise regression were determined by Bayesian information criterion (BIC).

## Results

### Identifying dietary manipulations that influence colitis severity

To identify macronutrients and individual ingredients that modulate experimental colitis, we used 32 unique diets (**Table S1** and **Table S2A**) with varied concentrations and sources of macronutrients comprising 31 ingredients whose nutritional content was dominated by a single macronutrient. For the macronutrients protein and fat, we tested a concentration range (wt/wt) of 5-41% and 1-21% respectively with soy protein, whey, casein, and egg whites as protein sources and corn oil, olive oil, lard, safflower oil, coconut oil, beef tallow, cotton seed oil, cocoa butter, palm oil, palm kernel oil, milk fat, and fish oil as fat sources. We separated carbohydrates into those that are largely host digestible, consisting of sucrose, fructose, glucose, maltodextrin, and corn starch tested over a range of 24-83%, and those that are largely host indigestible (fiber), consisting of cellulose, methylcellulose, psyllium, pectin, inulin, flaxseed, marshmallow root, potato starch, and slippery elm tested over a range of 5-15%. All diets were not limited for vitamins or minerals, and ranges for each macronutrient were selected to perturb each macronutrient concentration several fold.

Each dietary combination was fed *ad libitum* to 5-15 C56BL/6J SPF mice for one week, whereupon the animals were continued on the same diet while consuming 3% DSS in their drinking water to induce intestinal injury and inflammation (**Fig. 1A**). Importantly, unlike many colitis models, the DSS injury model is effective in both conventional mice and GF animals,^33–35^ allowing us to determine both the microbiota-dependent and microbiota-independent influences of diet on intestinal inflammation. During the initial dietary screen, disease severity was measured each day for seven days as the percent weight change of each animal relative to their mean baseline weight in the four days prior to DSS administration (**Table S2A**). Weight change was chosen as a metric of disease severity due to its ease of longitudinal measurement and its consistency with other inflammatory markers.^23^ We observed substantial variation in disease severity as a function of diet (p<0.0001, ANOVA).

**Figure 1.**
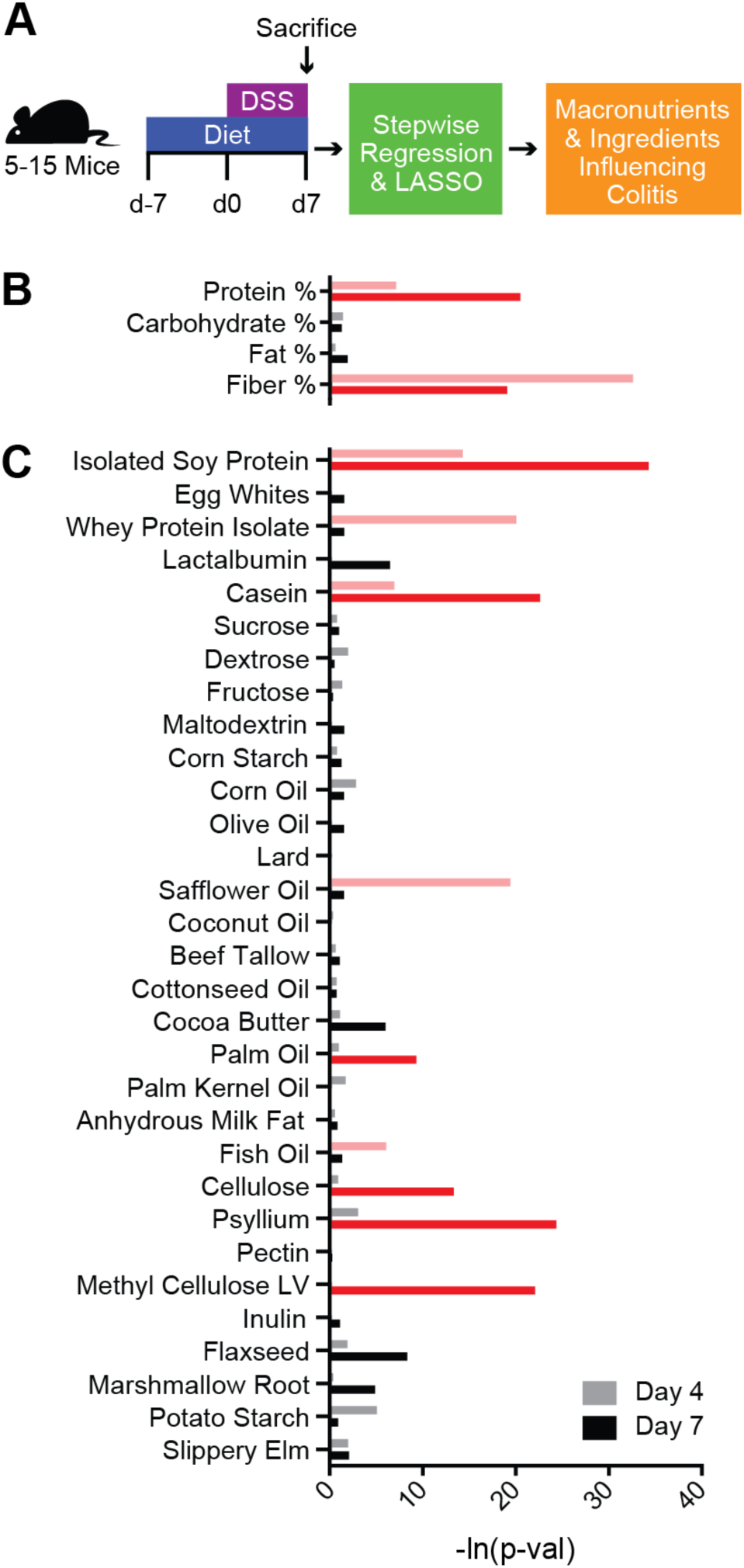
Dietary screening demonstrated that protein and fiber have the greatest effect upon DSS colitis. (A) Mice were fed one of thirty-two unique diets for a week before giving 3% DSS in drinking water. Disease severity was measured as weight loss. Stepwise regression (BIC selected factors highlighted in red) and lasso were used to identify (B) macronutrients and (C) dietary ingredients that contributed to differences in weight change at 4 and 7 days after beginning 3% DSS.

To understand the influence of macronutrients on disease severity, we modeled each animal’s weight loss during DSS administration as a function of the concentration of each macronutrient in the diet. We used stepwise regression and lasso^36^ to identify dietary macronutrients across the 32 dietary combinations that best explain the variation in disease severity estimated by weight loss at day 4 and at day 7 (**Fig. 1B**). Both stepwise regression and lasso identified protein as explaining increased weight loss at days 4 and 7 (p<0.001 for stepwise regression and lasso), while they identified fiber as explaining decreased weight loss at days 4 and 7 (stepwise regression; p=1.9×10^-14^ and p=9.5×10^-9^ respectively) or day 7 alone (lasso; p<0.0001) (**Table S3A**).

To understand the role of individual ingredients on the observed weight changes, we performed a regression analysis using the concentrations of the 31 different macronutrient sources used across the diets (**Fig. 1C**, **Table S3B**). Most protein sources (casein, isolated soy protein and egg whites) were identified as increasing weight loss, while psyllium fiber and pectin fiber were associated with decreased weight loss by lasso. Amongst the fats, palm oil and lard were associated with decreased weight loss by stepwise regression and lasso respectively, while fish oil was associated with increased weight loss on day 4 by stepwise regression.

We found both beneficial and deleterious influences of different fats on experimental colitis. However, most dietary fats had no effect or a slightly deleterious effect upon colitis. The association of fish oil with more severe disease supports a similar conclusion to previous work in an experimental colitis model,^37^ as well as recent clinical trials that found no benefit of omega-3 supplements in decreasing relapse in CD patients.^38^ To further explore the influence of fat on disease severity beyond the weight change based dietary screen, we compared DSS-treated mice fed high corn oil (20%, TD.09056; HCO) or low corn oil (1%; TD.09050, LCO) diets. Mice fed HCO and LCO experienced no significant difference in weight loss, colonic cytokines, or histopathology (**Fig. S1)**. The phenotypes significantly altered in response to changes in dietary fat concentration were fecal lipocalin-2 (p=0.024, t-test; **Fig. S1**) and colon length (p=0.0052, t-test; **Fig. S2**). Although the direction and scale of the host response to dietary fat was similar to previous studies (i.e., increased fat moderately exacerbates disease),^39,40^ the magnitude of the influence of dietary fat in DSS colitis was far less than either protein or fiber.

Across all diets and macronutrients, protein, in the form of casein, was identified as most significantly associated with increased weight loss while fiber, in the form of psyllium, was associated with the largest protection from disease. To explore these macronutrients with the greatest influence on disease severity, we focused on high casein (41%, TD.09054; HC) and low casein (6%, TD.09052; LC) diets, as well as diets containing psyllium (5%, TD.150229; PSY) or cellulose (5%, TD.09053; CEL). Mice fed a HC diet, while consuming 3% DSS, lost approximately twice as much weight by day 4 and 7 respectively as mice consuming LC diet (15.6% vs 8.2% on d4 and 35.8% vs 18.0% on d7; p<0.0001 both days, t-test; **Fig. 2A; Table S2A**). The HC-associated weight loss was associated with increased concentrations of fecal lipocalin-2 (p=0.0035, t-test; **Fig. 2C Table S4B**), increased TNF-α and IL-6 from colon explants (p=0.0003 and p=0.0033 respectively, t-test; **Fig. 2D-E; Table S4A**), more severe colon histopathology (p=0.0056, t-test; **Fig. 2F-G; Table S4C**), and decreased colon lengths (p=0.0003, t-test; **Fig. S2; Table S4D**). In contrast, mice fed PSY diet experience approximately 9-fold and 2-fold less weight loss on day 4 and 7 respectively when compared to mice fed CEL diet (0.8% vs 7.2% on d4 and 14.2% vs 23.8% on d7; p=0.010, p=0.011, t-test; **Fig 2B; Table S2A**). We also observed concomitant differences in fecal lipocalin-2 (p=0.0013, t-test; **Fig. 2C; Table S4B**), inflammatory cytokines from colon explants (TNF-α and IL-6; p=0.0077, p=0.0007, t-test; **Fig. 2D-E; Table S4A)**, colon histopathology (p<0.0001, t-test; **Fig. 2F-G; Table S4C**), and colon length (p=0.0003, t-test; **Fig. S2; Table S4D**). Together, these immune monitoring and pathology findings support the observations from our diet screen that changes in dietary protein and fiber are major determinants of disease outcome in the mouse DSS colitis model.

**Figure 2.**
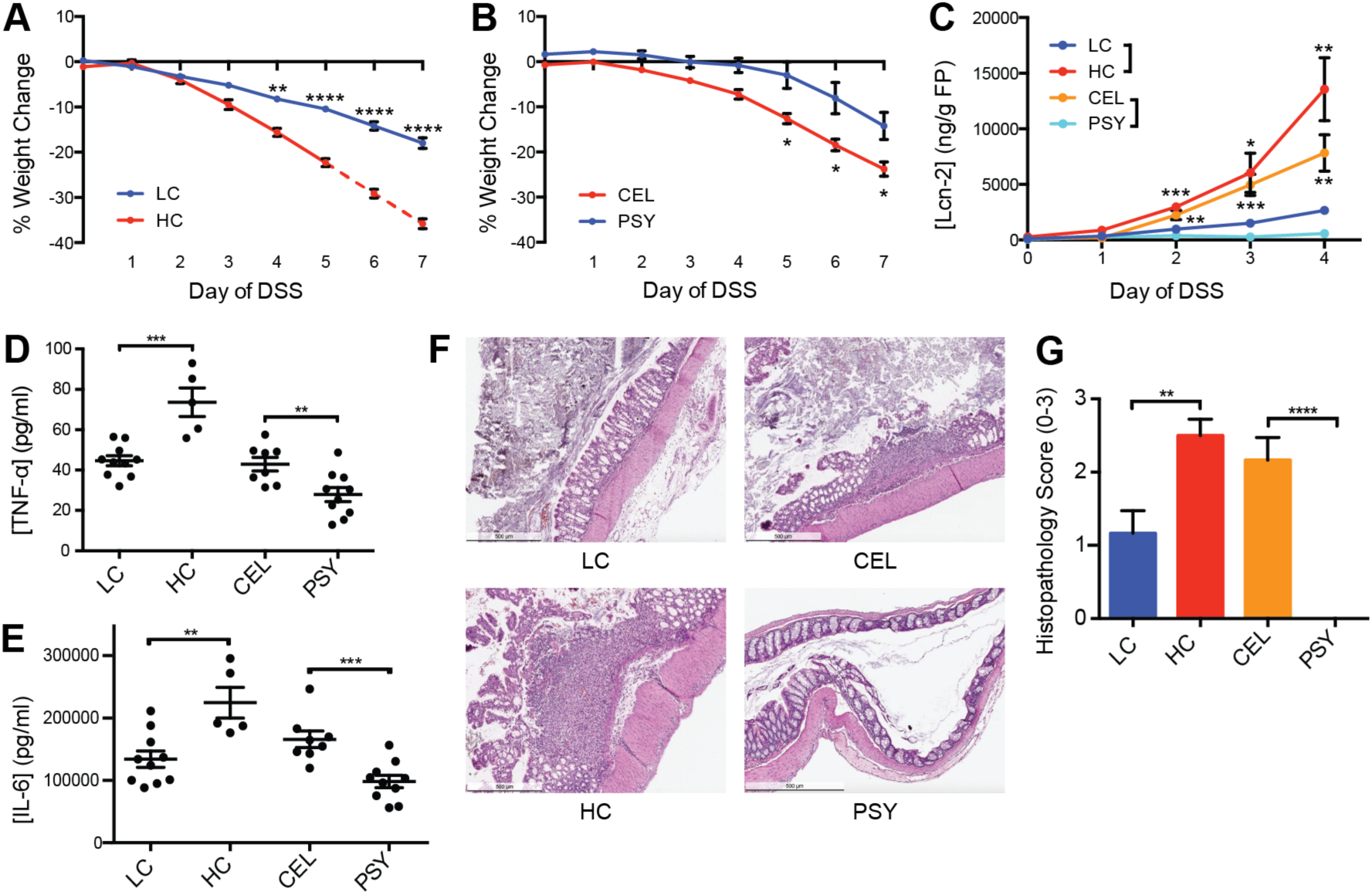
Casein promotes colitis and psyllium protects from colitis. Mice fed (A) LC diet had less severe weight loss after one week on DSS than those fed a HC diet, while mice consuming the (B) PSY diet had less severe disease than those consuming the CEL diet. (C) After 4 days of DSS, fecal lipocalin-2 was almost undetectable for the PSY diet and highly increased in the HC diet. After 7 days of DSS or at the humane endpoint, (D) TNF-α and (E) IL-6 from whole colonic explants demonstrated increases in HC and decreases in PSY. (F) Representative H&E staining and (G) histopathological scoring of colons indicate that HC and CEL drive more severe disease. The mean ± SEM are plotted for each time point and group (AC, G). Points represent individual animals (D-E). ^*^p<0.05,^**^p<0.01, ^***^p<0.001, ^****^p<0.0001

### Casein modulates colitis severity through microbiota-dependent mechanisms

To determine if the deleterious influence of protein on disease severity was dependent upon the microbiota, we provided HC or LC diet to both GF and SPF animals. Since GF mice have more severe disease than SPF in the DSS colitis model,^34^ we used 2% DSS in the GF group to induce equivalent disease to 3% DSS in the SPF mice (**Fig. S3**). We found a 1.8-fold increase (4.4 days to 7.8 days; **Fig. 3A**) in survival time following the start of DSS treatment for SPF mice on the LC diet compared with the HC diet (p<0.0001, Log-Rank test), while there was no difference in survival between mice consuming the LC and HC diets in GF animals (5.6/5.2 days, respectively; p=0.22, Log-Rank test), demonstrating that the influence of dietary casein concentration on disease severity is microbiota-dependent.

**Figure 3.**
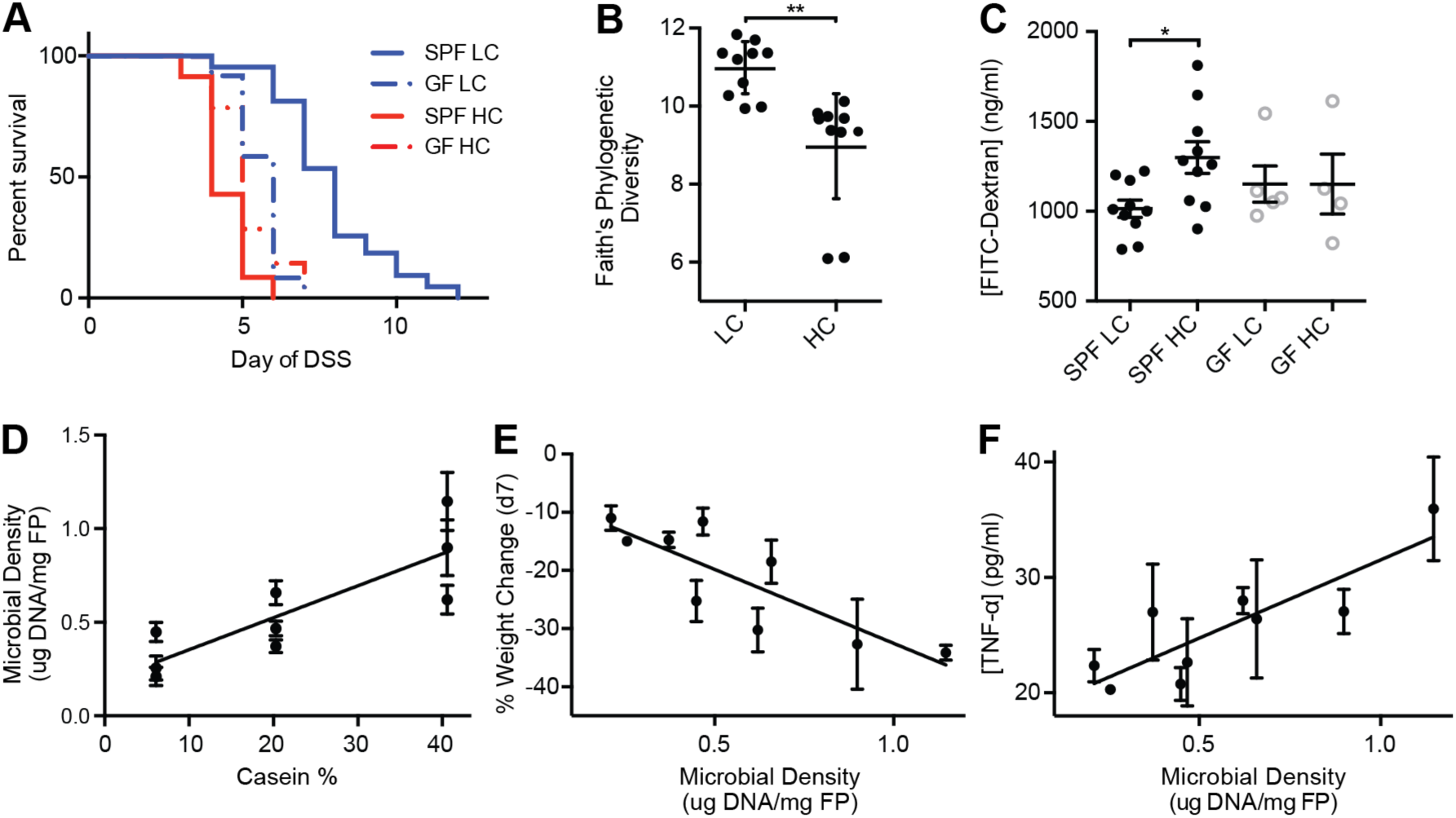
The effects of casein upon health and colitis are microbiota-dependent. (A) In the DSS colitis model, HC diet reduces survival relative to LC diet in SPF mice but not in GF animals. Alpha diversity measured by (B) Faith’s phylogenetic diversity rarified to 10k reads per sample demonstrated decreased microbial diversity in the HC diet. Furthermore, mice on HC diet had (C) decreased barrier function in healthy SPF mice, while barrier was not influence by casein in GF mice. In panes D-F, animals were given one of 9 controlled diets with different casein concentrations (TD.09049-TD.09057) with varying amounts of nutrients for a week and then given DSS. (D) Increased gut microbial density, driven by casein concentration, was associated with more severe disease in terms of both (E) weight change on DSS at d7 and (F) TNF-α. Lines represent surviving number of animals (A). Black (SPF) and gray (GF) points represent individual animals (B-C). The mean ± SEM are plotted for each point (D-F). *p<0.05, **p<0.01

To understand if casein might configure the microbiota to increase host susceptibility to DSS colitis, we analyzed the microbiota in animals fed HC and LC diets that were unchallenged with DSS. Culture independent 16S rDNA sequencing demonstrated a reduced diversity in the fecal microbiota of mice fed HC compared to LC diet (Faith’s phylogenetic diversity; p=0.0022, t-test; **Fig. 3B**). We also observed significantly different shifts in fecal microbiota composition between mice fed the LC and HC diets (weighted unifrac; p=0.001, permanova; **Fig. S4**), consisting of phylum-level increases in the relative abundance of Bacteriodetes and decreases the Firmicutes in the mice fed HC diet (**Fig. S4**). Although these particular changes in gut microbiota composition might modulate the microbiota-dependent influence of protein on colitis severity, we observed this influence of casein on colitis across independent experiments with differences in microbiota composition (**Fig. S4**), suggesting the reduction of diversity and increased susceptibility to DSS colitis is consistent across mouse different microbiotas.

We investigated features of unchallenged mice fed HC or LC diets that might explain the microbiota-dependent influence of protein on DSS colitis. In transcriptional profiles of colonic tissue from mice fed HC or LC diet, we found decreased casein associated with increased expression of genes involved in cell adhesion, such as AOCS3, LAMC1, KRT17, and DES, as well as increases in genes associated actin cytoskeleton and intermediate filament organization (**Fig. S5**). As we observed decreased expression of cell adhesion genes in mice fed a high casein diet, we hypothesized that dietary protein composition may affect the integrity of the intestinal barrier. By measuring translocation of orally administered FITC-dextran in the bloodstream of mice, we found mice fed HC diet had a microbiota-dependent increase in intestinal permeability compared to those consuming LC diet (**Fig. 3C; Table S6A**).

Previous studies have shown that antibiotics can reduce disease severity in DSS mouse colitis models^41^ and provide some benefit in patients with IBD.^42^ Importantly, work in both rat and mice DSS models suggest the reduction in colitis severity upon antibiotic treatment is proportional to the ability of the antibiotic to deplete the microbiota.^43^ We previously identified casein as a potential driver of gut microbial density.^11^ We therefore hypothesized that increased microbial density, associated with high casein diets relative to low casein diets, could drive the diet and microbiota dependent changes in pathology. We analyzed the microbiota density of the 9 different diets, from our initial screen (TD.09049-TD.09057; 5-10 mice per diet, **Table S2A** see green highlights), that contained various amounts of casein (low, medium, and high), corn oil (low, medium, and high) and sucrose (low, medium, and high). As expected in mice unchallenged with DSS, increasing casein was significantly correlated with an increase in gut microbial density (R^2^=0.71, p=0.0045, F-test; **Fig. 3D**; **Table S2A**) with a 3-fold average difference in microbial density between the HC and LC diets.

To determine if increases in gut microbial density might create an environment for elevated colitis susceptibility, we measured the gut microbiota density in the same mice after administration of 3% DSS and found increased dietary protein also increased gut microbial density under inflammatory conditions (R^2^=0.88, p=0.0002, F-test; **Fig. S6**; **Table S2A**). Protein-driven changes in gut microbial density were significantly associated with disease severity estimated as weight change on day 7 of DSS administration as well as with colonic TNF-α and IL-10 from colon explants (R^2^=0.59, 0.73, 0.45; p=0.016, 0.0035, 0.049 respectively, F-test; **Fig. 3E-F**, **Fig. S6**, and **Table S2A**). These results suggest the level of inflammation in DSS injury is driven in part by the density of bacteria within the intestinal lumen, which would also explain the microbiota-dependent nature of the influence of casein on disease severity. To validate that casein-driven increases in microbial density are causally linked to increased pathology in the DSS model, we fed groups of mice the LC diet, the HC diet, or the HC diet supplemented with the antibiotic metronidazole, which induced a similar gut microbial density to the LC diet (**Fig. S7**). We found no significant difference in survival between mice receiving the HC + metronidazole or the LC diet when administered 3% DSS (p=0.53, Log-Rank test; **Fig. S7**). Both of these microbial-density lowering interventions enabled significantly longer survival than seen in mice fed HC diet (p=0.0039, p=0.0033, respectively, Log-Rank test; **Fig. S7**). Together these results show that the microbiota-dependent increase in DSS pathology in animals fed HC diet compared to LC diet is linked to increased microbial density. This increase in microbial density in HC diet, which also correlates with a reduced microbial diversity and decreased intestinal barrier function, might exacerbate DSS induced intestinal injury either through increases in antigenic load that elevate the likelihood of microbial invasion into the nearby tissue or increased capacity to generate pathogenic metabolites.

### Psyllium modulates intestinal permeability and colitis severity

To determine if psyllium, the most protective ingredient in our dietary screen, mitigated disease in a microbiota-dependent manner, we measured survival of SPF and GF mice fed either a high psyllium diet (15% psyllium, TD.130633; HPSY) or high cellulose diet (15% cellulose, TD.130630; HCEL) for a week prior to and during induction of colitis. SPF mice fed the HPSY diet survived on average for 27.6 days, 4.6 times longer than mice fed the HCEL diet (6 days, p<0.0001, Log-Rank test; **Fig. 4A**). In contrast to the microbiota-dependent effects of casein, GF animals experienced some of the protective benefit from consuming HPSY diet and survived for an average of 16.7 days, 3.6 times longer than GF mice fed HCEL diet (4.7 days, p<0.0001, Log-Rank test; **Fig. 4A**). Although psyllium provided some microbiota-independent benefit in the context of DSS-colitis, we observe significantly greater benefit in SPF animals on the HPSY diet relative to GF animals (p<0.0001; Log-Rank test; **Fig. 4A**) suggesting that some of the benefit is also microbiota-dependent.

**Figure 4.**
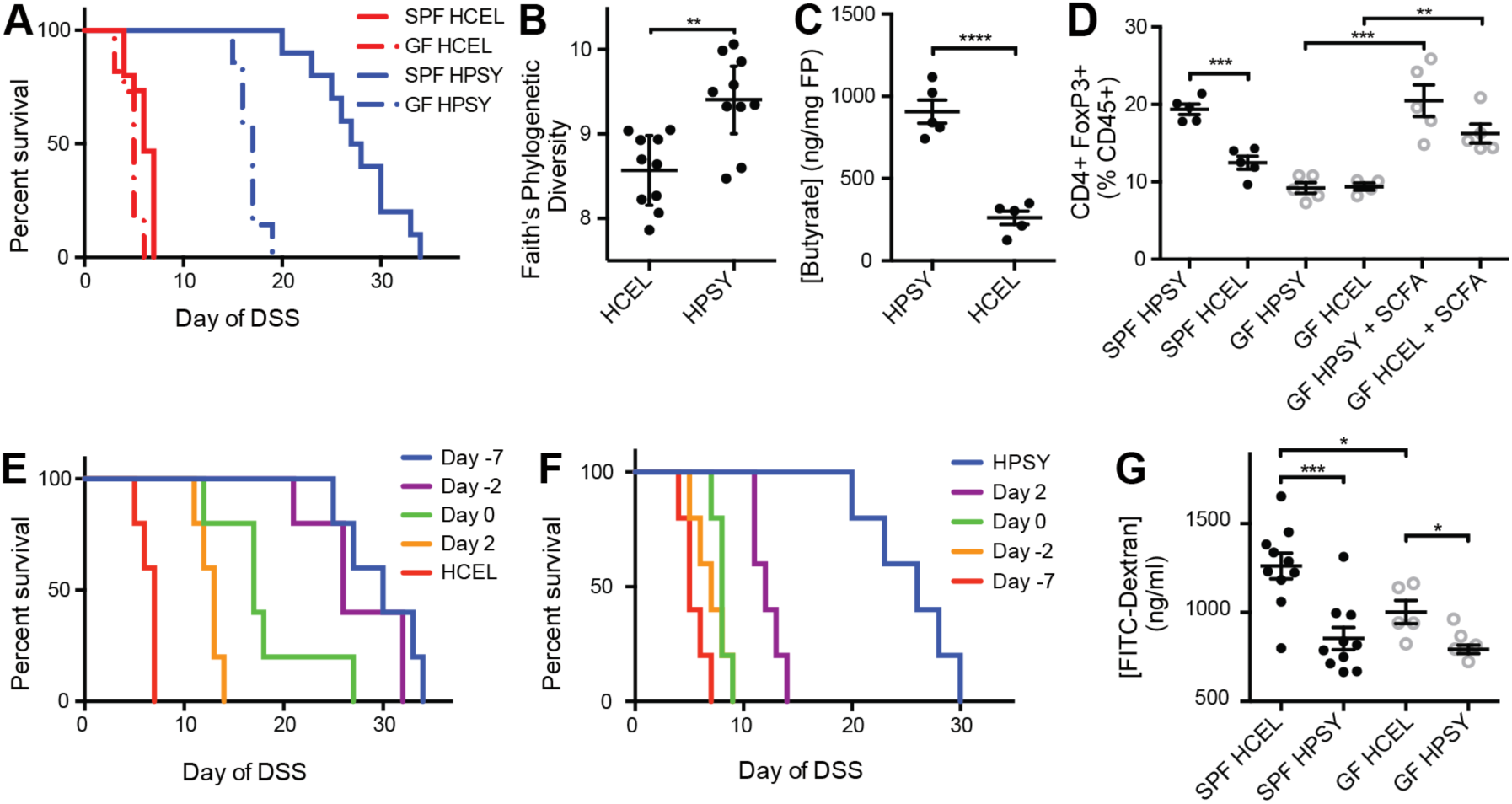
Dietary psyllium has both microbiota-dependent and microbiota-independent effects upon host health and colitis. The HPSY diet led to (A) increased survival in both SPF and GF mice with the greatest difference observed in SPF animals. Alpha diversity measured by (B) Faith’s phylogenetic diversity rarified to 7k reads per sample demonstrated increased microbial diversity in the HPSY diet. Mice fed HPSY had increased colonic (C) butyrate, isobutyrate (p=0.028, t-test; **Table S5**), and valerate (p<0.0001, t-test; **Table S5**), which was coupled with (D) an increase in colonic Tregs in SPF animals. In GF animals, Tregs were increased via oral supplementation of SCFA in the drinking water. (E) To determine the dietary pretreatment time necessary to obtain the protective benefit of psyllium in DSS colitis, mice were initially given the HCEL diet and were transferred to the HPSY diet at various time points. Administering the HPSY diet at least two days prior to the administration of DSS was sufficient to provide the same protection as a week of diet. (F) To determine the duration of the benefit of psyllium in DSS colitis, mice were initially given the HPSY diet and were switched to the HCEL diet at different time points. The replacement of psyllium with cellulose almost immediately halted the protective benefits of psyllium in mouse survival. (G) The HCEL diet was associated with increased intestinal permeability with a larger barrier defect in SPF relative to GF mice. Lines represent surviving number of animals (A, E-F). Black (SPF) and grey (GF) points represent individual animals (B-D, G).^**^p<0.01, ^***^p<0.001, ^****^p<0.0001.

To better understand the potential role of the microbiota in the beneficial properties of psyllium dietary fiber, we performed 16S rDNA sequencing of fecal pellets taken after a week of HPSY or HCEL diet in mice unchallenged with DSS. The fecal microbiota in mice fed HPSY was more diverse than those fed HCEL (Faith’s phylogenetic diversity, rarefied to 7K reads; p=0.0012, t-test; **Fig. 4B**) and formed a distinct cluster from the gut microbiota of animals consuming the HCEL diet (weighted unifrac, rarefied to 7K reads; p=0.006, permanova; **Fig. S8**). Independent batches of mice with distinct microbiotas displayed similar responses to the same dietary intervention with increased psyllium driving increased microbiota diversity in unchallenged animals and less severe disease in those animals upon administration of DSS.

Certain dietary fibers, like psyllium, provide a substrate for bacterial fermentation that generates SCFA, which increase the number and function of CD4+FoxP3+ regulatory T cells (Tregs) and have a protective effect in mouse colitis models.^28,44,45^ We therefore hypothesized that psyllium minimized disease in DSS colitis in a microbiota-dependent manner through increased production of SCFA and Tregs. Replacing dietary cellulose with psyllium led to increased concentrations of butyrate in the intestinal lumen (p<0.0001, t-test; **Fig. 4C**, **Table S5**), increased Tregs cells (p=0.0002, t-test; **Fig. 4D**), and less severe colitis in the T cell transfer model of colitis (**Fig. S9**). To determine if increased Tregs were responsible for the microbiota-dependent protective effects of psyllium in the DSS model, we directly increased Tregs in GF mice through oral administration of SCFA.^28^ However, exogenous SCFA were insufficient to protect GF mice in the DSS model (p=0.31, Log-Rank test; **Fig. S10**), suggesting that psyllium-driven, microbiota-dependent increases in SCFA and increased Tregs could not explain the microbiota-dependent protective effects of psyllium in the context of DSS colitis.

To test the kinetics of dietary psyllium protection, we fed mice the HCEL diet and then replaced it with the HPSY diet at various time points before and after beginning DSS treatment. We observed similar protective effects when we provided HPSY diet at either 2 or 7 days prior to DSS treatment (4.3/4.7 fold increased survival; p=0.0034, p=0.0030, respectively, Log-Rank test; **Fig. 4E**) with significant but less pronounced effects when psyllium was provided at the at the start of DSS administration or thereafter. To determine the duration of protection after psyllium consumption was halted, we replaced HPSY diet with HCEL diet at different time points before or after beginning DSS treatment. Animals given psyllium for the duration of the experiment had the longest survival (4.7 fold increased survival; p=0.0018, Log-Rank test; **Fig. 4F)** with a rapid decrease in benefit when the HPSY diet is removed (2.3 fold increased survival for d2; p=0.0045, Log-Rank test). Together these results demonstrate that the beneficial effects of psyllium are rapid and short-lived, which argues against an important role for the adaptive immune system and in favor of more rapid responses, like innate immune mechanisms, altered barrier function, or transcriptional changes.

To test the impact of psyllium on barrier function, we measured intestinal permeability with FITC-dextran in unchallenged SPF and GF mice fed the HPSY or HCEL diet. In both SPF and GF animals, mice fed HCEL had higher intestinal permeability than HPSY (p=0.0005, p=0.017 respectively, t-test; **Fig. 4G; Table S6A**). However, as in the protein dietary interventions, there was a microbiota-dependent difference in the influence of dietary fiber on intestinal permeability with decreased barrier function in SPF consuming HCEL relative to GF mice (p=0.040, t-test; **Fig. 4G; Table S6A**). To better understand the mechanism of the increased barrier function, transcriptional profiling was performed on colonic tissue (**Fig. S5**). We found HPSY diet drove changes in expression of genes related to the innate immune response (i.e., neutrophils and macrophages) and related to responses to bacteria/LPS (e.g., NCF2, PLSCR4, and TIMP4).

Overall these results indicate the benefits of psyllium in reducing DSS pathology are rapid acting and short-lived and unlikely to result from changes to the adaptive immune system. In unchallenged mice, microbiota-independent and microbiota-dependent benefits of psyllium are associated with improved intestinal barrier and increased gene expression related to innate immunity and responses to bacteria. These results suggest psyllium is strengthening antimicrobial immunity and limiting access of microbe invasion into the host tissue with perhaps some benefit resulting from the increased diversity of the microbial community associated with psyllium.

### The combinatorial influence of casein and psyllium on host physiology and inflammation

A critical feature of human diets is the continuous variation and combinations of multiple ingredients that, in concert, might alter disease risk or pathology. To quantify the combined influence of the two ingredients with the most significant impact on disease variation, we generated diets with all nine possible combinations of low, medium, and high casein (6, 20, 41%) and psyllium (0, 0.5, 5%; **Table S1** and **S2B**). We modeled weight change following DSS administration as the combined influence of both ingredients. We observed a significant influence of both psyllium and casein on weight change, as the beneficial influence of increased psyllium could offset the damaging influence of increased casein. The two extremes of high casein with low psyllium and low casein with high psyllium had the most and least severe disease respectively (**Fig. 5A**). To quantify the individual and synergistic influence of each ingredient on weight change, we performed a linear regression with each ingredient separately and combined. A linear regression with casein alone explains 33% of the variation in disease (estimated as weight change at DSS day 7; p=3.47×10^-5^; F-test) compared to 31% for psyllium alone (p=7.41×10^-5^, F-test) and 62% for casein and psyllium together (p=4.64×10^-10^, F-test; best model by BIC). Similar to our analyses of the effects of casein and psyllium individually, we also found a combinatorial influence of these two dietary ingredients on host physiology and the microbiota in unchallenged mice for both gut microbial density (R^2^=0.46, p=2.80×10^-6^, F-test) and intestinal permeability estimated with FITC-dextran translocation (R^2^=0.63, p=2.41×10^-4^, F-test; **Fig. 5B; Table S6B**). The combination of gut microbial density and intestinal permeability in unchallenged mice also explain most variation in DSS disease severity (71%; p=2.61×10^-5^, F-test). Altogether, these results indicate the influence of diet on colitis is the result of the combined influence of multiple ingredients and that studying any individual ingredient in isolation can be confounded by the contributions of other dietary factors.

**Figure 5.**
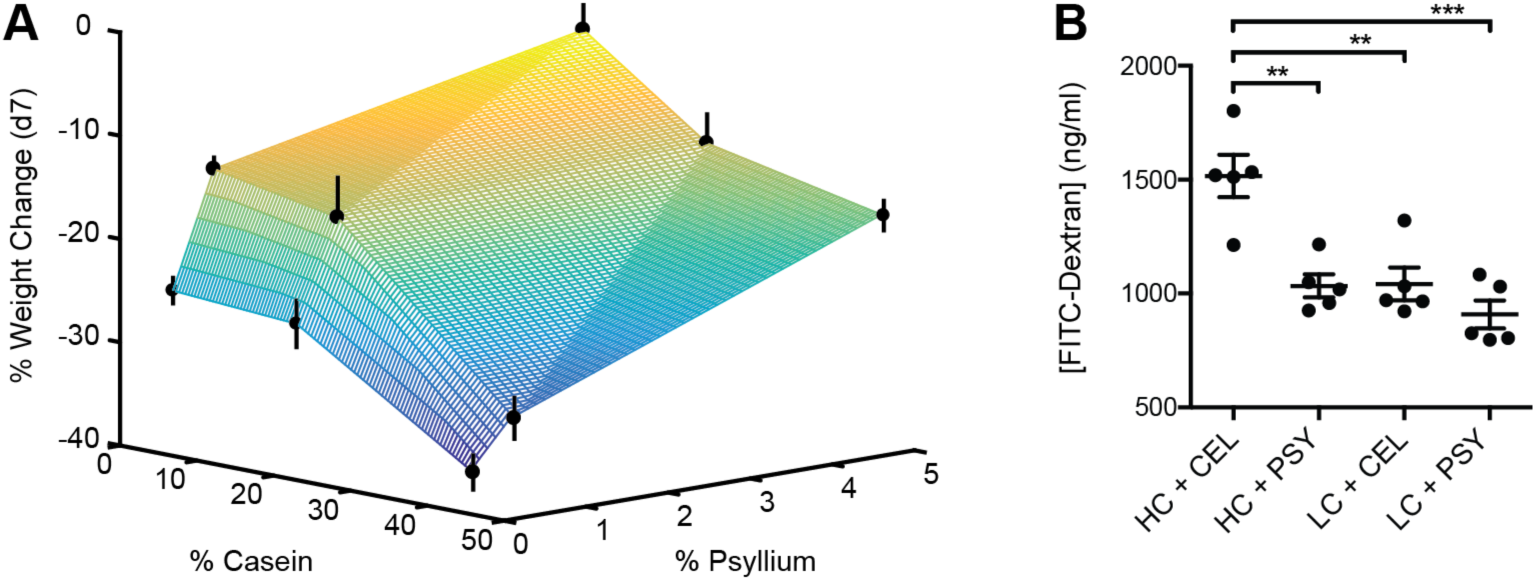
Dietary ingredients have combinatorial effects that alter host health and colitis severity. To examine the combinatorial interactions of casein and psyllium on host health and disease, nine diets (TD.09052-TD.09054, TD.140768-TD.140769, TD.150226-TD.150229) with various concentrations of casein (6%, 20%, 41%) and psyllium (0%, 0.5%, and 5%) were given to SPF mice a week prior to DSS. (A) Disease severity (i.e., weight change on DSS at d7) indicated the deleterious influence of increased dietary casein could be offset by increases in beneficial psyllium fiber. The most severe colitis is associated with high casein and no psyllium fiber, while mice consuming low casein and higher psyllium fiber are largely protected from colitis. These dietary ingredients also had a combinatorial effect on host physiology in mice unchallenged with DSS. Mice given the four extremes of the nine diet matrix (i.e., combinations of high/low casein and psyllium) showed similar patterns in (B) barrier function—high casein decreasing barrier function could be offset by adding psyllium. The mean ± SEM are plotted for each group (A) and black points represent individual SPF animals (B). The colored surface is a triangulation-based linear interpolation of the data (A). ^**^<0.01, ^***^p<0.001

## Discussion

Deciphering the complex interactions between the gut microbiota, diet, and IBD remains a major challenge in the development of diet-based therapies for the prevention and treatment of IBD. Towards this, we used mouse models of colitis to control host genotype and the gut microbiota, while systematically manipulating dietary composition. In our study of over 40 dietary combinations tested in the DSS intestinal injury model of colitis, we observed significant microbiota-dependent and microbiota-independent influences of diet on experimental colitis, as well as on host physiology in the context of healthy, unchallenged animals. Overall, we observed a remarkable 6.3-fold difference in survival from the most deleterious (4.4 days survival) to the most beneficial diet (27.6 days survival) in the DSS model, highlighting the importance of diet in experimental colitis. As in previous studies,^39,40^ we identified a small but significant influence of dietary fat on experimental colitis severity. By exploring all macronutrients and multiple macronutrient sources in the same dietary screen, we determined that the impact of dietary fat on colitis is far less than dietary protein or fiber.

With the exception of methylcellulose fiber that appeared to exacerbate disease similar to a previous study on carboxymethylcellulose,^17^ most dietary fibers were either beneficial or had little to no effect on colitis. Recent results have highlighted the important role of fiber in maintaining intestinal homeostasis.^19,20^ Our results further support the protective role of certain fibers both in their ability to increase barrier function in unchallenged animals as well as their ability to minimize disease in the DSS injury model of colitis and the adoptive T cell transfer model of colitis. Although EEN provides a clinically validated diet-based therapy for IBD,^1^ many enteral nutrition formulas are notably lacking in fiber. Moreover fiber is excluded from the diet of some IBD patients due to its potential to lead to blockages in the context of stricturing disease.^2^ However, a recent dietary survey of over 1500 individuals with IBD found the avoidance of fiber is associated with a greater risk of a flare in CD.^46^ Psyllium was the most effective fiber in slowing experimental colitis progression. In a placebo-controlled trial, psyllium was associated with a significantly higher rate of improvement in gastrointestinal symptoms in UC in remission compared to a non-gel-forming placebo fiber.^47^ In our DSS colitis experiments, the amount of psyllium sufficient for small benefits was extremely low (i.e., 0.5% (wt/wt)) with increased benefits with dosage escalations up to 15% suggesting a benefit might be possible with minimal dietary supplementation (**Fig. 5A** and **Fig. S11**).

In addition to providing further evidence of the importance of dietary fiber in mitigating intestinal inflammation, we found a novel microbiota-dependent interaction between dietary protein, gut microbial density, and intestinal inflammation whereby increases of protein drive an expansion of the gut microbiota leading to an exacerbation of disease in the DSS colitis model. As seen in small intestine bacterial overgrowth syndrome, increases in microbial density within the gut can lead to IBD-like symptoms.^48^ The important role observed for protein supports the observation in a prospective cohort study that found consumption of meat and protein were associated with increased likelihood of UC relapse.^49^

Our data demonstrate that the influence of diet on gut inflammation involves numerous potentially additive or offsetting effects between dietary components and the gut microbiota that should be considered when developing specific IBD diets in the clinic. Dietary trials that exclude specific proinflammatory ingredients will only be successful in extreme cases, as the majority of diet’s influence on gut inflammation is likely the sum of the effect of multiple dietary components - balancing these components represents the key to effective diet therapy. As we gain a broader knowledge of the role of dietary components in IBD, we might develop boundary conditions and constraints to enable consumption of a diverse palette of foods while simultaneously avoiding combinations or quantities of dietary components that exacerbate disease. Finally, it is important to note our observation that dietary macronutrients that alter disease severity also influence host physiology in healthy, unchallenged animals, suggesting that diet could prevent disease relapse for patients who already have IBD or even prevent the occurrence of the disease in individuals at risk for developing IBD.

## Abbreviations used in this paper

IBD: Inflammatory Bowel Disease
CD: Crohn’s Disease
UC: ulcerative colitis
EEN: exclusive enteral nutrition
DSS: dextran sodium sulfate
HC: high casein diet
LC: low casein diet
PSY: psyllium diet
CEL: cellulose diet
HPSY: high psyllium diet
HCEL: high cellulose diet
SCFA: short chain fatty acids
SPF: specific pathogen free
GF: germ free
Tregs: T regulatory cells

## Acknowledgements

We are grateful to C. Fermin, E. Vazquez, R. Ng, and G. Escano in the Mount Sinai Immunology Institute Gnotobiotic facility for their help with gnotobiotic animal husbandry. C. Berin, C. Yang, O. Vennaro, and B. Mickelson provided helpful suggestions during the course of this work. Metabolite measurements were performed by the Stable Isotope and Metabolomics Core Facility of the Diabetes Research and Training Center (DRTC) of the Albert Einstein College of Medicine supported by NIH/NCI grant P60DK020541. Next generation sequencing was performed at NYU School of Medicine by the Genome Technology Center partially supported by the Cancer Center Support Grant, P30CA016087.

## Funding

This work was supported by grants from the NIH (NIGMS GM108505, NCCIH AT008661, and NIDDK DK108487) and SUCCESS.

## Author contributions

S.R.L. and J.J.F. designed the experiments; S.R.L., A.M., and G.J.B. generated the data involving immune function and inflammation; E.J.C. developed high throughput methods for measuring gut microbial density; S.R.L., G.B., A.G., E.J.C, J.F.C., J.C.C., and J.J.F. analyzed the data; S.R.L. and J.J.F. wrote the paper. In addition to supplemental tables, raw data suitable for computational modeling are provided as supplemental data.

## SUPPLEMENTARY FIGUERS

**Figure S1.**
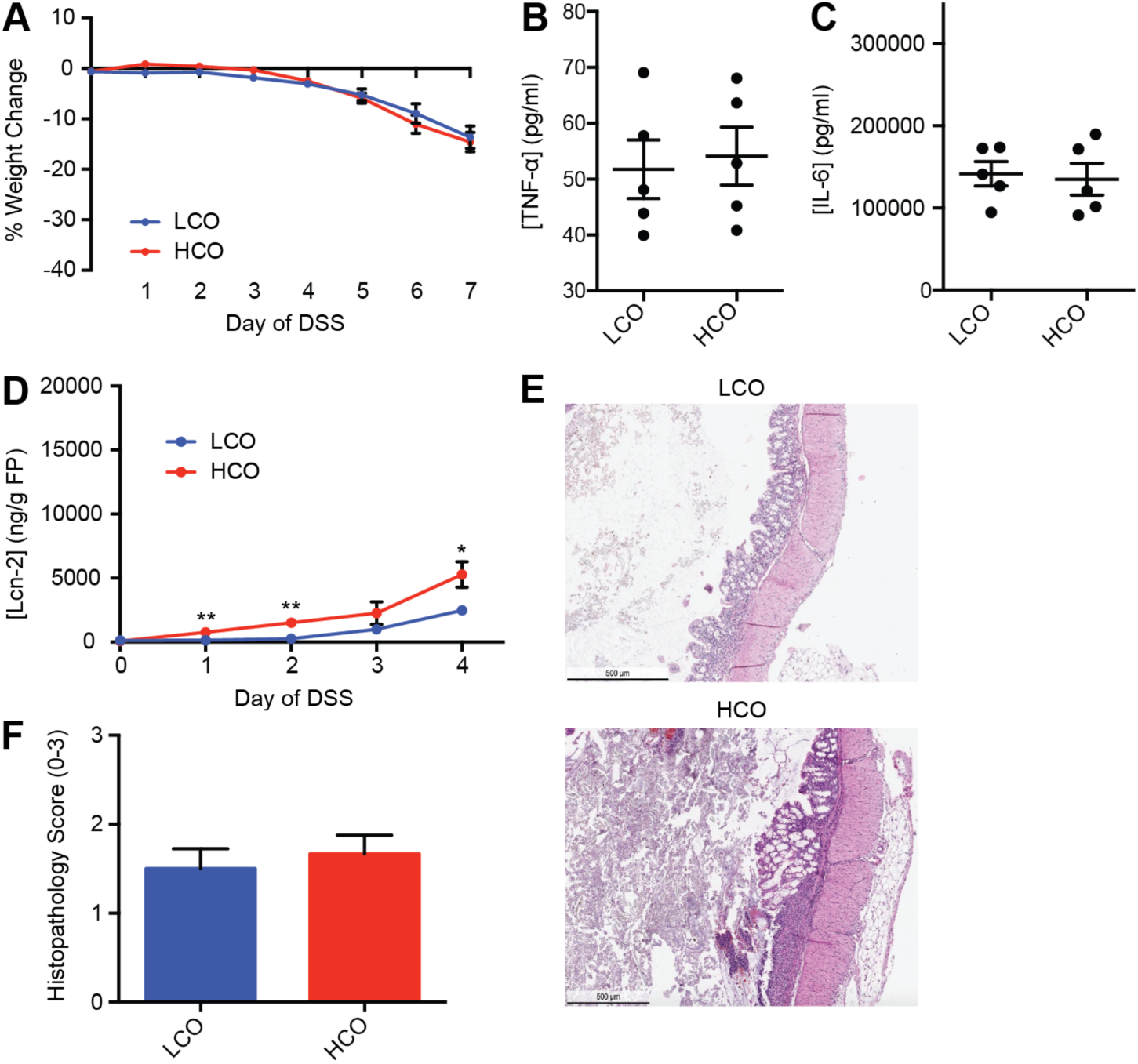
Dietary fat has a minor impact on DSS colitis severity. Animals were fed LCO or HCO diet for a week followed by the same diet with DSS for 7 days (panes A-C) or 4 days (panes D-F). As a measure of disease severity, (A) weight change, colonic cytokines (B) TNF-α, and (C) IL-6, (D) fecal lipocalin-2, and (E) histopathological scoring and (F) representative H&E staining were used. Percent weight change, TNF-α, IL-6, and histopathological scoring indicated no significant differences in disease severity. Fecal lipocalin-2 and colon length (**Fig. S1**) were significantly altered by dietary fat. The mean ± SEM are plotted for each time point and group (A,D,F), while black points represent individual animals (B-C). ^*^p<0.05, ^**^p<0.01

**Figure S2.**
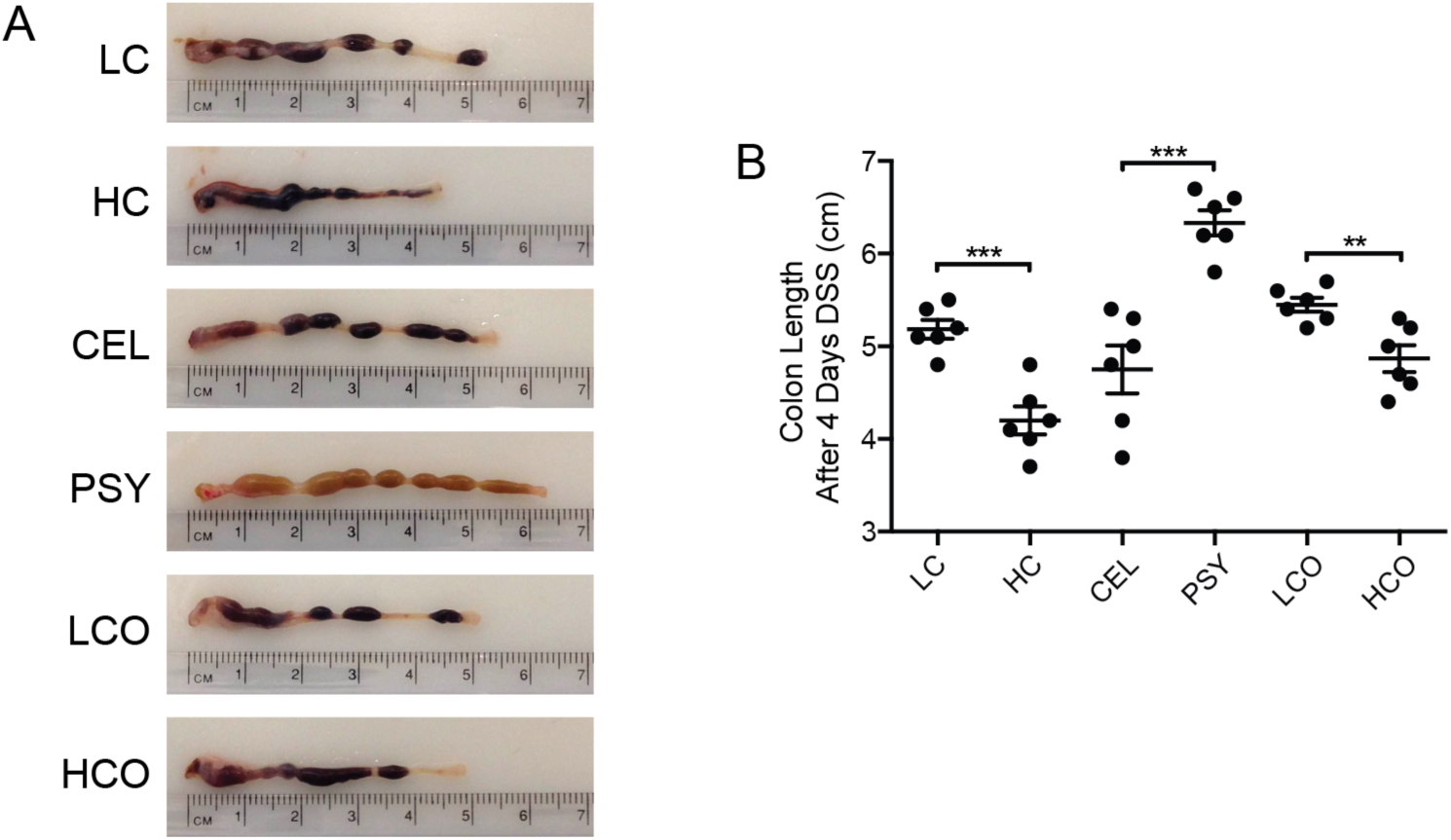
Diet alters colon lengths and colonic tissue in DSS colitis. Mice were given one of six diets (LC, HC, CEL, PSY, LCO, and HCO) for a week, followed by the same diet and DSS for four days. (A) Representative colons and (B) the combined analysis of all colon lengths indicate that colon lengths vary by diet in a similar manner to other markers of intestinal inflammation (e.g., **Fig. 2** and **Fig.S1**). As expected, the largest differences were between the HC and LC diets and between the CEL and PSY diets. Points represent individual animals. Mean ± SEM are plotted. ^**^p<0.01, ^***^p<0.001

**Figure S3.**
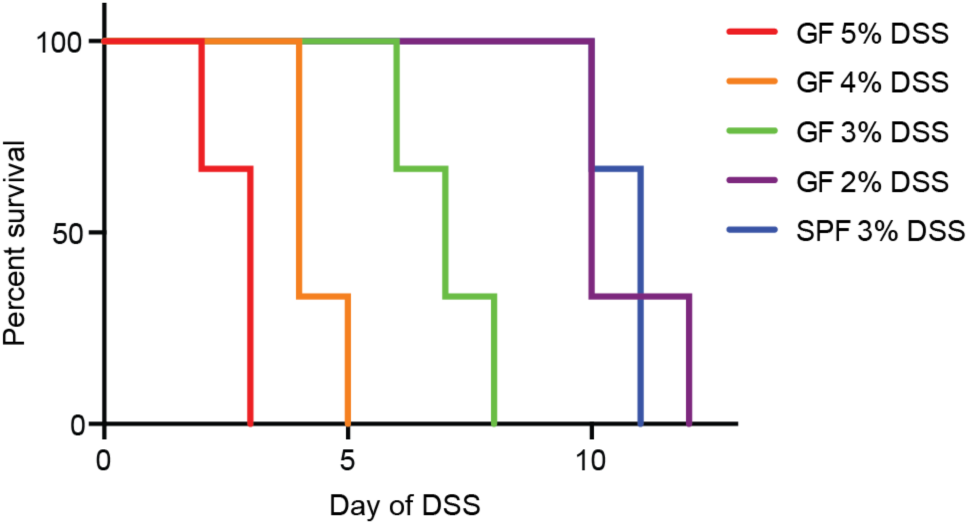
DSS titration survival curves in germ-free animals. DSS is more potent in GF mice compared to SPF mice. GF mice were fed standard chow (TD.2108S) and given 2-5% DSS. 2% DSS in GF mice (purple) leads to equivalent survival to 3% DSS in SPF mice (blue).

**Figure S4.**
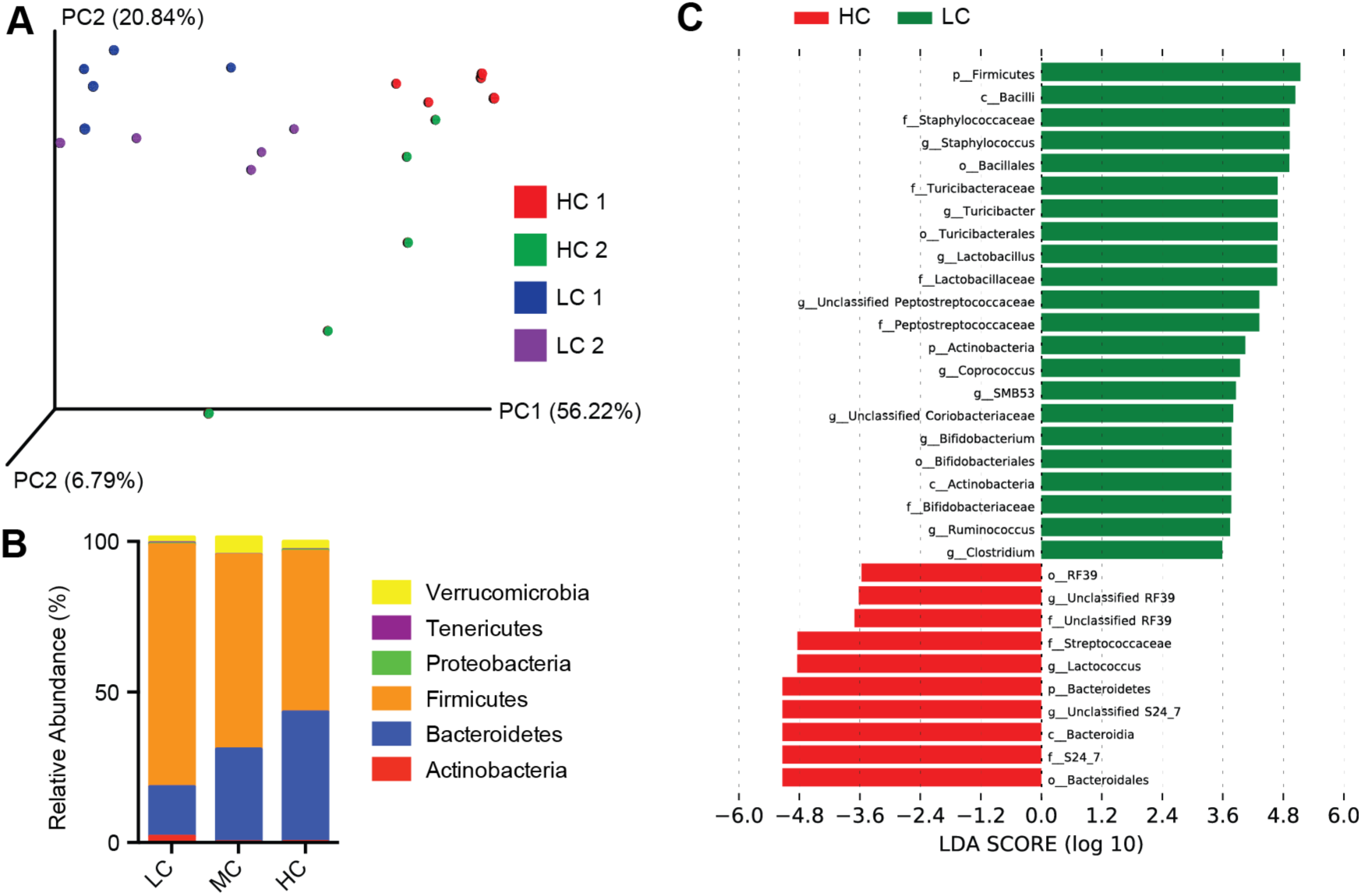
Changes in the gut microbiota in response to differences in dietary casein. 16S rDNA amplicon sequencing of the V4 hypervariable region was performed on fecal samples collected from animals consuming a HC, MC (medium casein, 20%, TD.09053), or LC diet. Beta diversity was measured by (A) weighted UNIFRAC at 10K reads and visualized by PCOA. Across two separate experiments, LC animals and HC animals clustered together and were significantly different from each other (p=0.001, permanova) from each other. There was also a grouping based on the mouse batch (1 or 2) suggesting similar community changes occur in response to dietary protein manipulation in two different microbiotas. (B) At the phylum level, increasing casein increases the relative abundance of Bacteriodetes and decreases the Firmicutes. (C) LEfSe analysis revealed phylum Firmicutes, class Bacilli, family Staphylococcaceae, genus Staphylococcus, and order Bacillales as the groups of bacteria increased the most in LC, whereas bacteria from order Bacteroidales, family S24_7, class Bacteroidia, Genus Unclassified S24_7, and phylum Bacteroidetes as the groups of bacteria increased in HC.

**Figure S5.**
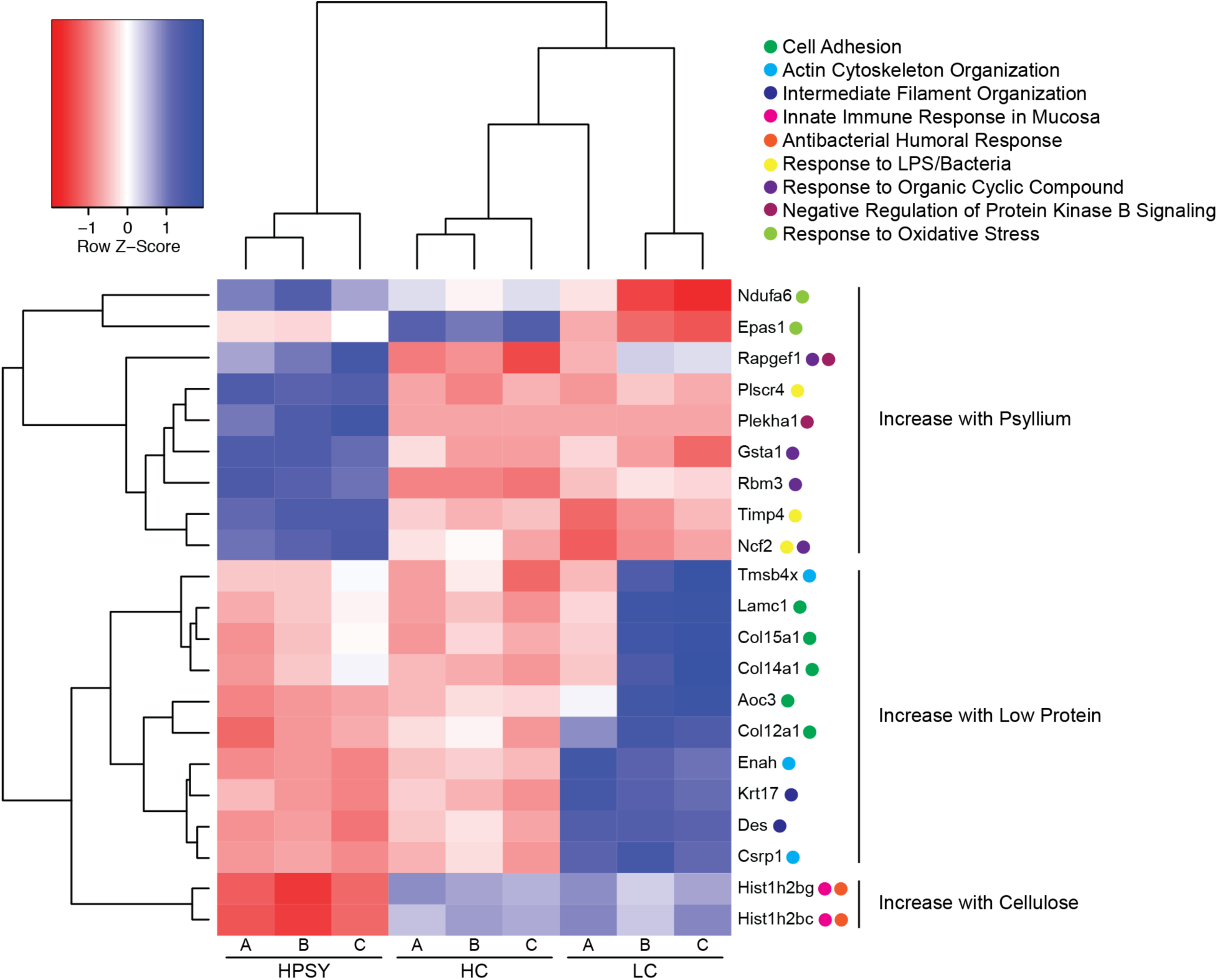
mRNA transcription changes to dietary modulation. To determine the effects of diet upon host transcription, we performed RNA-Seq on mice given one of three diets (HPSY, HC, LC) for a week prior to tissue collection. Heatmap shows raw z-scores clustered by gene and diet. Gene ontology enrichment analysis was performed and biological function annotations were given for each gene with DAVID bioinformatics resources.^50^

**Figure S6.**
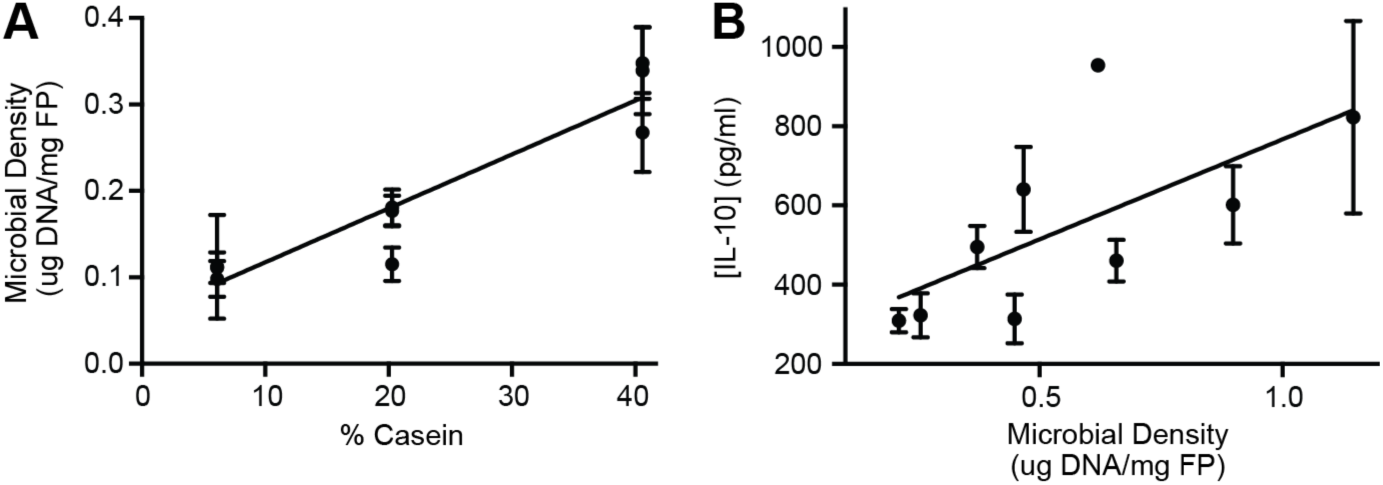
Casein is a major determinate of gut microbial density and disease severity in DSS colitis. Increasing casein increases the gut microbial density (**Fig. 3D**). To determine the effects of DSS upon gut microbial density, we gave SPF mice different casein concentrations from 9 controlled diets (TD.09049-TD.09057) for a week prior to giving DSS. After two days of DSS, (A) gut microbial density was quantified. Similar to healthy mice, mice on DSS had increased gut microbial density with increased casein concentration (R^2^=0.88, p=0.0002, F-test); however, giving DSS had an ~3-fold decrease in gut microbial density compared to healthy animals. Similar to TNF-α (**Fig. 3F)**, (B) IL-10 increases as gut microbial density increased (R^2^=0.45, p=0.049, F-test).

**Figure S7.**
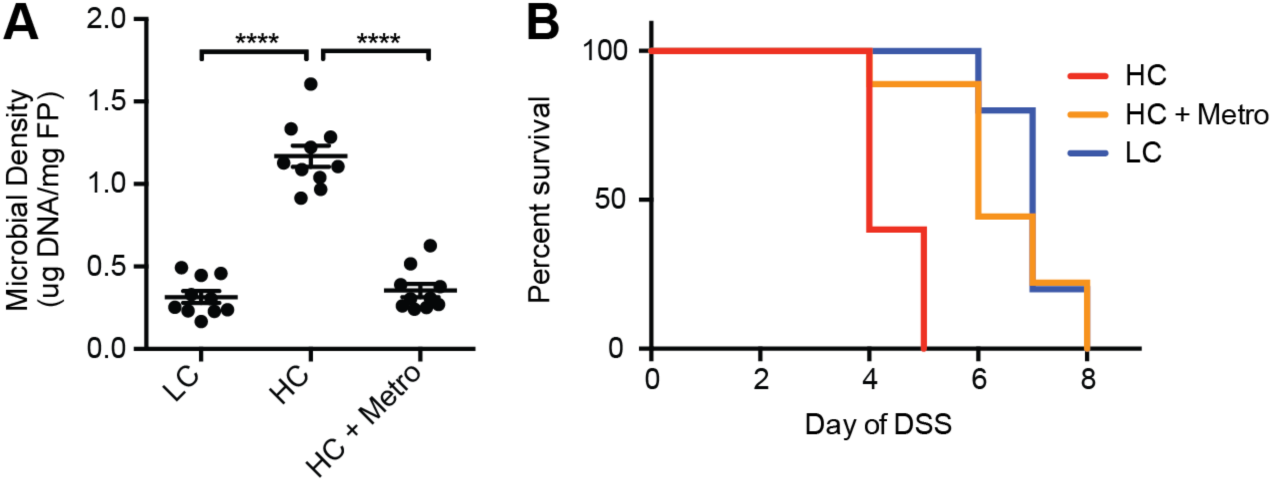
Metronidazole rescues the high casein phenotype by lowering gut microbial density. Mice were given LC diet, HC diet or HC diet + metronidazole (HC + Metro) with similar (A) gut microbial density values between LC diet and HC diet + metronidazole after being on diet + antibiotics for 7 days. (B) Mice in each of the groups were given DSS, which showed antibiotic-based depletion of the gut microbiota density to levels similar to LC diet could increase survival.

**Figure S8.**
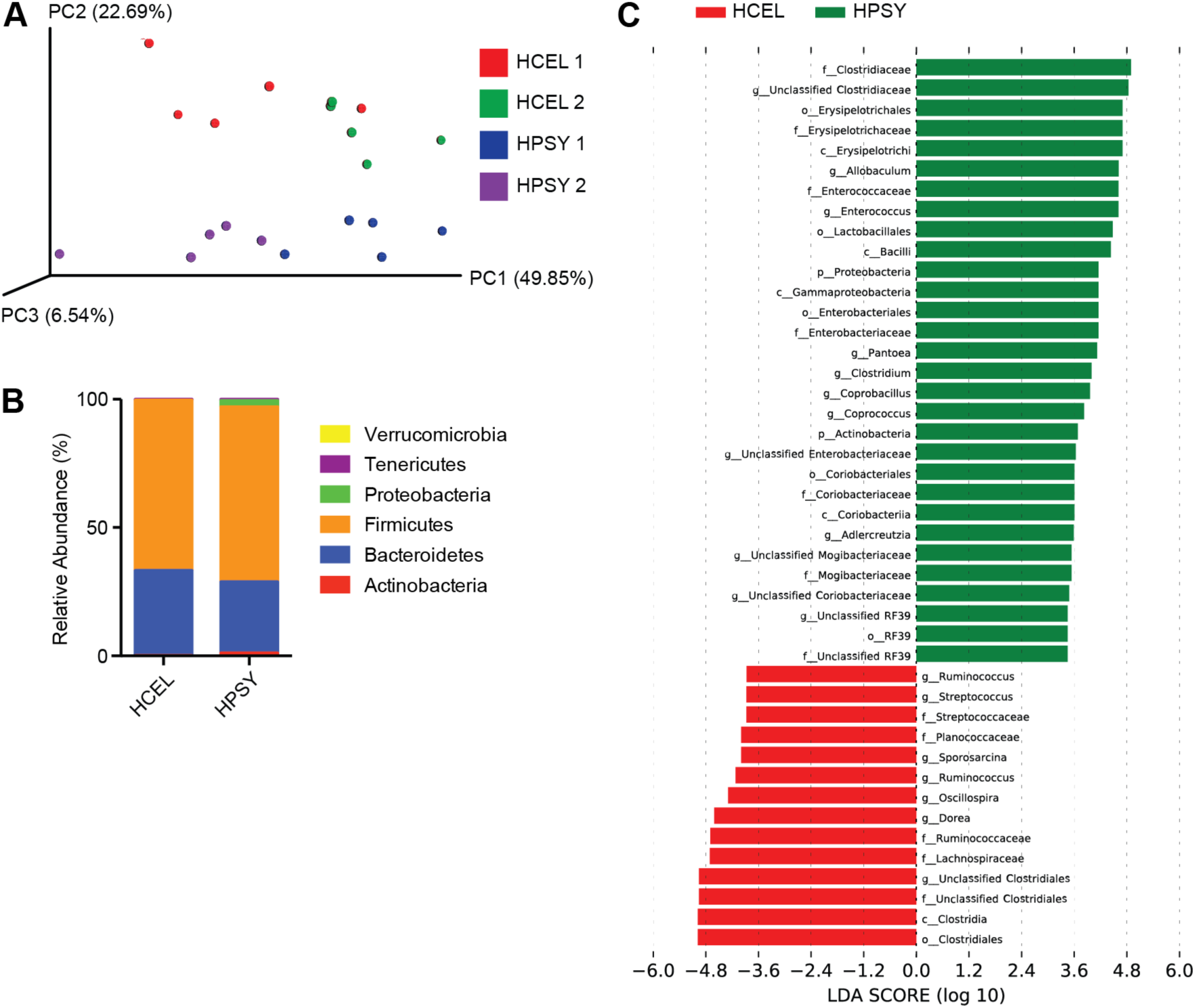
Alterations of the microbiota associated with perturbations to dietary fiber. We performed 16S rDNA amplicon sequencing of the V4 hypervariable region on fecal samples from two independent groups of mice consuming HPSY or HCEL diet. Beta diversity was measured by (A) weighted UNIFRAC at 7K reads and visualized by PCOA. Across two separate experiments, microbial communities clustered by diet and were significantly different from each other (p=0.006, permanova). There was also a grouping based on the mouse batch (1 or 2) suggesting similar community changes occur in response to dietary protein manipulation in two different microbiotas. (B) There were no major differences in relative abundances at the phylum level. (C) LEfSe analysis demonstrated that HPSY increased the relative proportions of bacteria from the family Clostridiaceae, genus Unclassified Clostridiaceae, order Erysipelotrichales, family Erysipelotrichaceae, and class Erysipelotrichi, while the bacteria from the order Clostridiales, class Clostridia, family Unclassified Clostridiales, genus Unclassified Clostridiales, and family Lachnospiraceae had the greatest growth in the HCEL diet.

**Figure S9.**
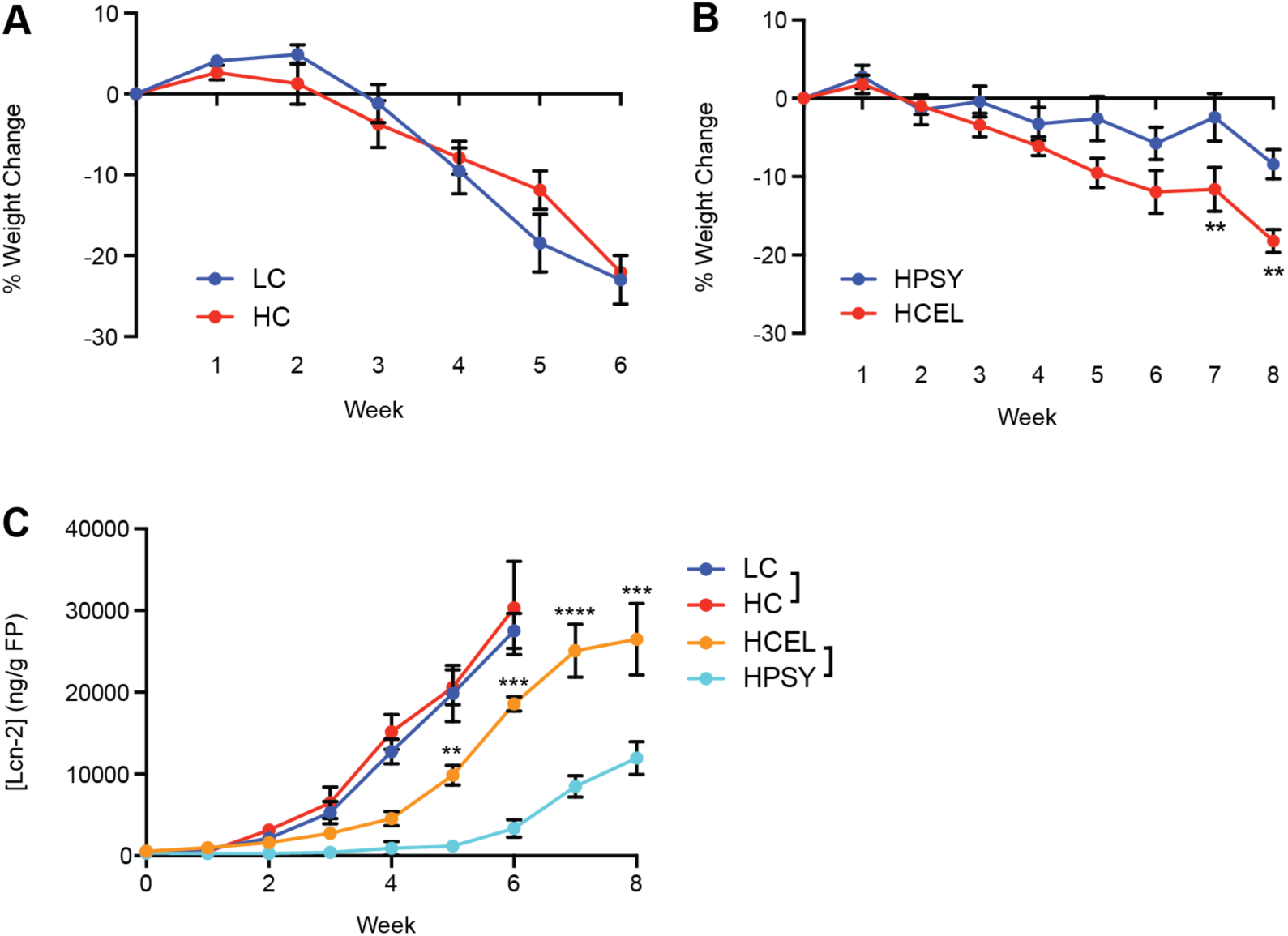
Effects of casein and psyllium in the adoptive T cell transfer colitis model. Starting a week prior to T cell transfer animals were fed HC, LC, HPSY, or HCEL diet for the duration of the experiment. (A) There were no differences in weight loss between mice given the HC and LC diets, suggesting the significant influence of protein on colitis is not T cell driven. (B) Mice consuming HPSY had less severe disease, measured by (B) weight loss and (C) fecal lipocalin-2. The protective benefit of HPSY suggested that both the microbiota-dependent and microbiota-independent effects of psyllium are protective in both the T cell transfer and DSS colitis models. The mean ± SEM are plotted for each time point. ^**^p<0.01, ^***^p<0.001, ^****^p<0.0001

**Figure S10.**
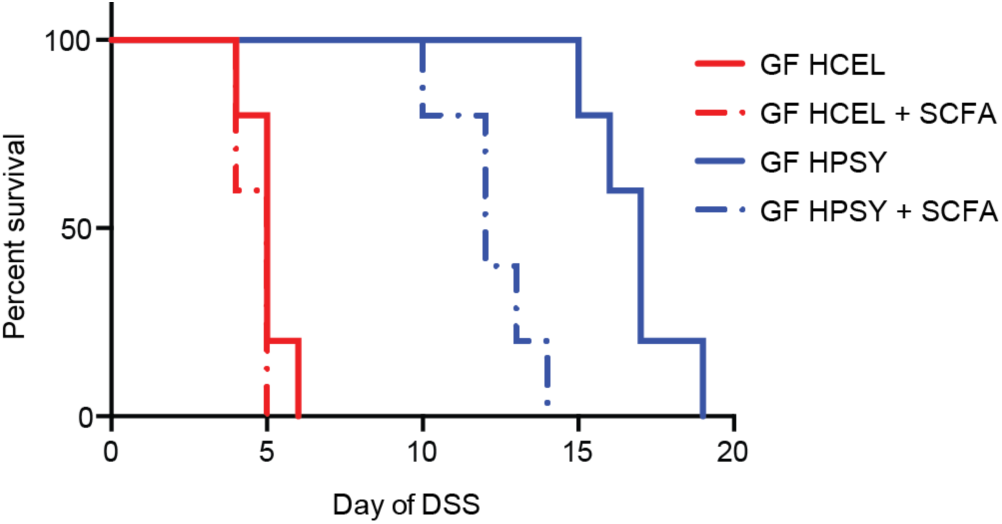
Survival curves in the GF DSS colitis model with diet ± SCFA supplementation. GF animals were given either the HPSY or HCEL diet ± SCFA (40mM butyrate, 25.9mM propionate, 67.5mM acetate in drinking water) for 4 weeks before DSS was given. In healthy GF animals given SCFA for 4 weeks, Tregs were increased (**Fig. 4D**).^28^ Survival was measured as time until a humane endpoint. HPSY, but not SCFA supplementation, significantly increased survival (p<0.0001, Log-Rank test).

**Figure S11.**
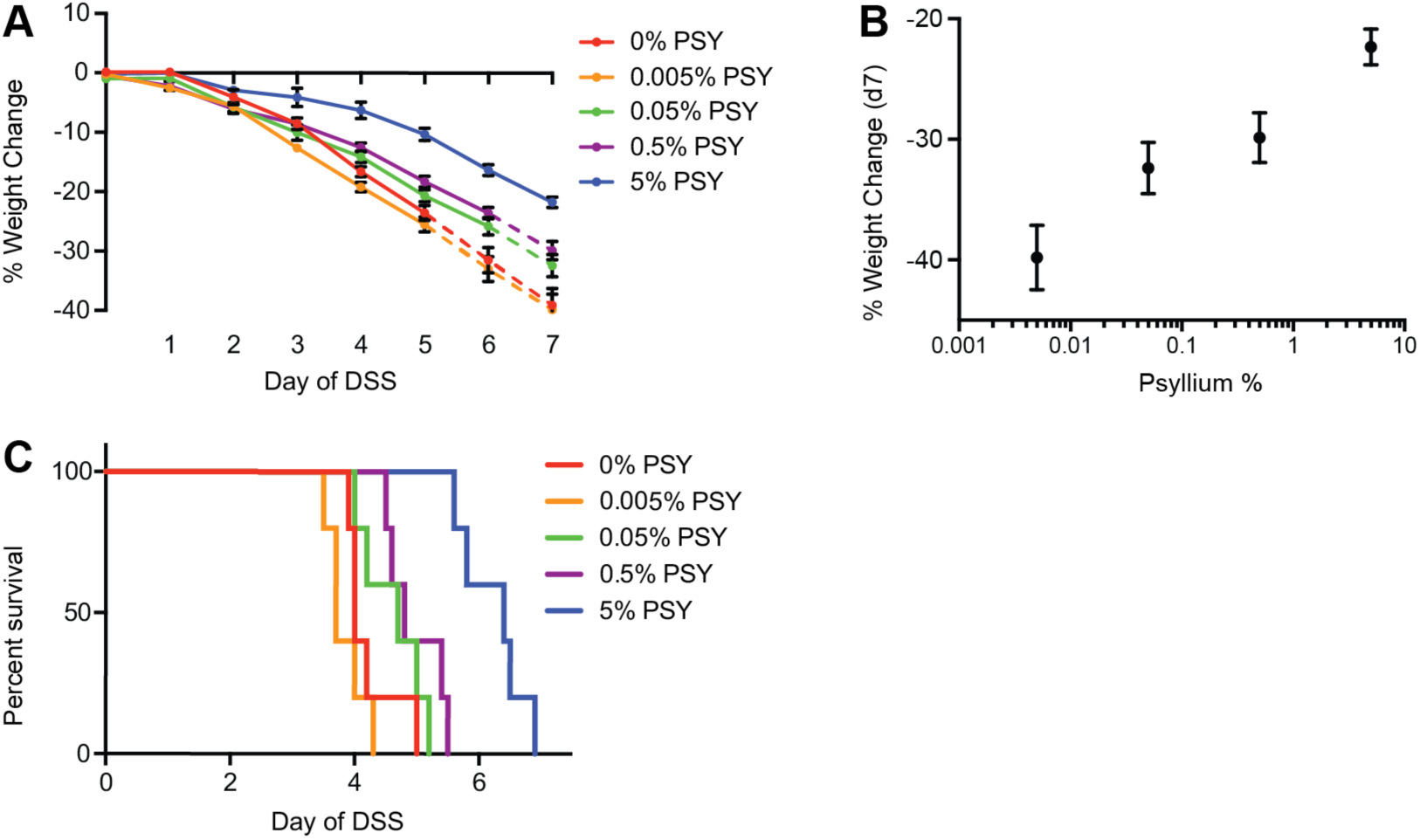
Psyllium dose response in DSS colitis. To determine the effective beneficial range of psyllium fiber, we developed four trial diets with high casein (41%) and concentrations of psyllium from 0.005%-5% (0.005% psyllium, TD.140771; 0.05% psyllium, TD.140770; 0.5% psyllium, TD.140769; 5% psyllium, TD.140768). Each diet was provided to mice for a week before providing DSS. Disease severity was measured as (A) percent weight loss over time, (B) percent weight change on DSS at day 7, and (C) survival. Psyllium provided a benefit at concentrations as low as 0.5% (p=0.022, t-test, day 7 weight loss; p=0.060, Log-Rank, survival), with increased benefit at increased psyllium concentrations. The mean ± SEM are plotted for each time point.

**Figure S12.**
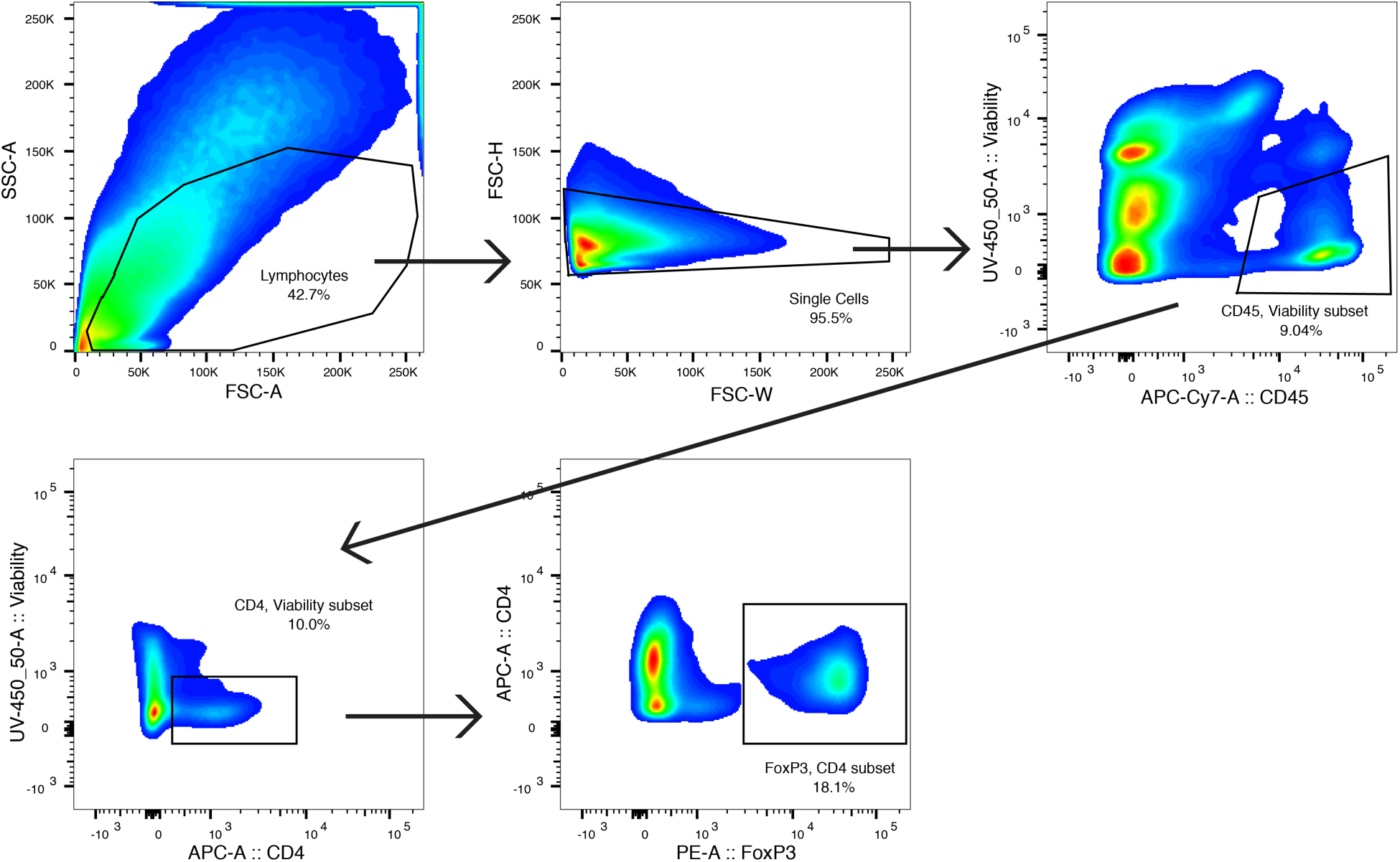
Gating scheme for determining the populations of CD4+FoxP3+ Tregs. Colonic lamina propria lymphocytes were identified using FSC-A and SSC-A. From that population, single cells were selected by FSC-W and FSC-H. Viable CD45+ cells were gated to remove dead cells and non-hematopoietic cells and a gate for viable CD4+ cells was used as an additional filter. Tregs were identified by CD4+FoxP3+ staining.

